# Local Attachment Explains Small-World-Like Properties of Fibroblastic Reticular Cell Networks in Lymph Nodes

**DOI:** 10.1101/369801

**Authors:** Kasper M.W. Soekarjo, Johannes Textor, Rob J. de Boer

## Abstract

Fibroblastic reticular cells (FRCs) form a cellular network that serves as the structural backbone of lymph nodes and facilitates lymphocyte migration. This FRC network has been found to have small-world properties. Using a model based on geographical preferential attachment, we simulated the formation of a variety of cellular networks and show that similar small-world properties robustly emerge under such natural conditions. By estimating the parameters of this model, we generated FRC network representations with realistic topological properties. We found that these properties change markedly when the network is expanded from a thin slice to a 3D cube. Typical small-world properties were found to persist as network size was increased. The simulated networks were very similar to 2D and 3D lattice networks. According to the used metrics, these lattice networks also have small-world properties, indicating that lattice-likeness is sufficient to become classified as a small-world network. Our results explain why FRC networks have small-world properties and provide a framework for simulating realistic FRC networks.

## Introduction

Lymph nodes play a central role in immune responses by functioning as a “meeting point” for lymphocytes. The structural framework of lymph nodes is a reticular network that consists of elastin and reticular fibers. These fibers are made of collagen and are surrounded by fibroblastic reticular cells (FRCs) [1]. The FRC network is formed through FRCs establishing connections with each other and forming inter-cellular channels [2], induced by signals from lymphocytes [3]. Besides having structural purposes, the FRC network facilitates lymphocyte migration within the lymph node [4–6].

In order to gain a deeper understanding of the properties and organization of the FRC network, several imaging techniques have been used [1,7,8]. The topology of the FRC network was recently reconstructed by modeling the connections found in confocal microscopy images of a cross-section of a lymph node [9]. In order to obtain network properties that accurately represent the observed FRC networks, this was done for several lymph nodes obtained from different mice. Methods from graph theory were employed to analyze the properties of the network.

Within network theory, an important property is the “small-worldness” of a network. A small-world network combines strong local connectivity, measured by a high clustering coefficient, and strong global connectivity, measured by a short path length between any two nodes in the network. The high clustering of small-world networks leads to high robustness against random damage to the network [10]. It should be noted that a small-world network is not necessarily scale-free, which is a network type in which the degree distribution follows a power law [11] and which is also highly robust to random damage [12]. Small-world properties have been detected in a multitude of biological networks, such as gene expression [13] and metabolism [14], and in a wide variety of other real-world networks including railways [15], the airport network [16] and communication networks [17]. By quantifying the topology of reconstructed FRC networks mathematically, Novkovic et al. found that the network exhibits small-world properties, and suggested that these might have evolved under evolutionary pressure [9].

Geographic attachment models are capable of generating small-world networks [18, 19]. These models connect pairs of nodes based on the physical distance between them, with a closer proximity resulting in a higher probability of forming a connection. Longer connections arise when connected nodes separate due to network expansion. Because FRCs are expected to preferentially connect to nearby FRCs during the development of the FRC network, we here study whether a similar method suffices for the small-world topology described by Novkovic et al [9]. Our model is capable of reliably generating random networks with properties similar to those of the observed FRC network. Small-world properties were found to emerge naturally and robustly, indicating that these properties are inherent to networks of cells that tend to connect to nearby cells. Furthermore, two- and three-dimensional lattice networks were found to exhibit similar small-world properties, demonstrating that a lattice-like structure can be sufficient to be classified as a small-world network.

## Methods

### Measuring small-worldness

Watts and Strogatz originally introduced small-world networks as a combination of the path length of random networks (which grows logarithmically with the number of nodes in the network) and the high clustering coefficient of a lattice [20]. This definition is often used as the key criterion for models generating small-world networks [18, 19, 21, 22], but it can only be used when networks with multiple sizes are available to calculate the growth of the average shortest path length. This measure is therefore not applicable to many real-world networks for which only a single size is available.

In order to quantify small-worldness in single size networks, several metrics have been developed based on comparing the clustering coefficient and path length of the observed network to that of a random or lattice network with the same number of nodes and edges [23–26]. We employ the same metrics to measure small-worldness as Novkovic et al. [9] used to compare generated networks with observed FRC networks. The two parameters used to assess small-worldness are *σ* and *ω* defined as follows:

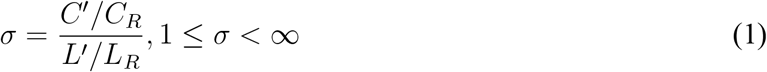

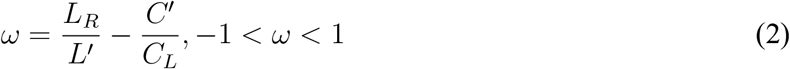

where *C′* and *L′* are the average clustering coefficient and the average shortest path length of the network, *C_R_* and *L_R_* are the average clustering coefficient and the average shortest path length of an equivalent random network, and *C_L_* is the clustering coefficient of an equivalent lattice network. Equivalent networks are formed with the same amount of nodes and edges using either a random or a 1D ring lattice structure respectively. A network is considered to be small-world if *σ* > 1 and −.5 > *ω* > 0.5 [23–25]. Furthermore, a network is classified as lattice-like if *ω* < 0 [25]. It is well known that *σ* grows with the size of the network [24], and hence *ω* is typically the most important measure.

### Simulation of FRC network

The observed network by Novkovic et al. originated from a 300×300×30 μm cross-section of the T-cell zone of a mouse lymph node [9]. This cross-section is scaled to a 1×1×0.1 volume in our model. Each FRC is considered to be a single node and is placed uniformly at random in the volume. The average diameter of a FRC has been observed to be approximately 10 μm [3, 9] (which is scaled to a diameter of 0.03 for each node). Nodes are replaced to a new random position when they overlap with other nodes. Generation and analysis of networks was done using the NetworkX package in Python (code is available in supplementary materials).

### Generating connections between cells

In the model, the probability of any two nodes forming a connection is determined through a probability function parameterized by the distance between those nodes. First, the FRC network consists of physical connections between cells, suggesting that there should be a maximal length for connections. Second, long connections are assumed to be increasingly difficult to achieve, and to occur less frequently than short connections. We implement these principles by using a monotonically decreasing probability function with a maximal length threshold:

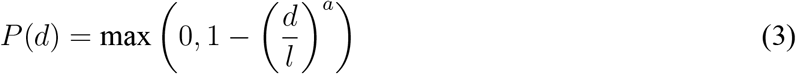

This function gives the probability *P* (*d*) for two nodes at distance d forming a connection. The probability of connecting is greater than zero when the distance is smaller than the maximal length threshold l (Fig. 1). The probability function decreases from 1 to 0 on the domain 0 ≤ *d* ≤ *l* and the shape of the function is given by the parameter a, with *a* > 0 (Fig. 1). Using a we can vary the ratio between long and short connections. Higher values of a result in a slower decrease of the probability to connect and therefore increase the fraction of long connections in the network (Fig. 1).

**Figure 1:**
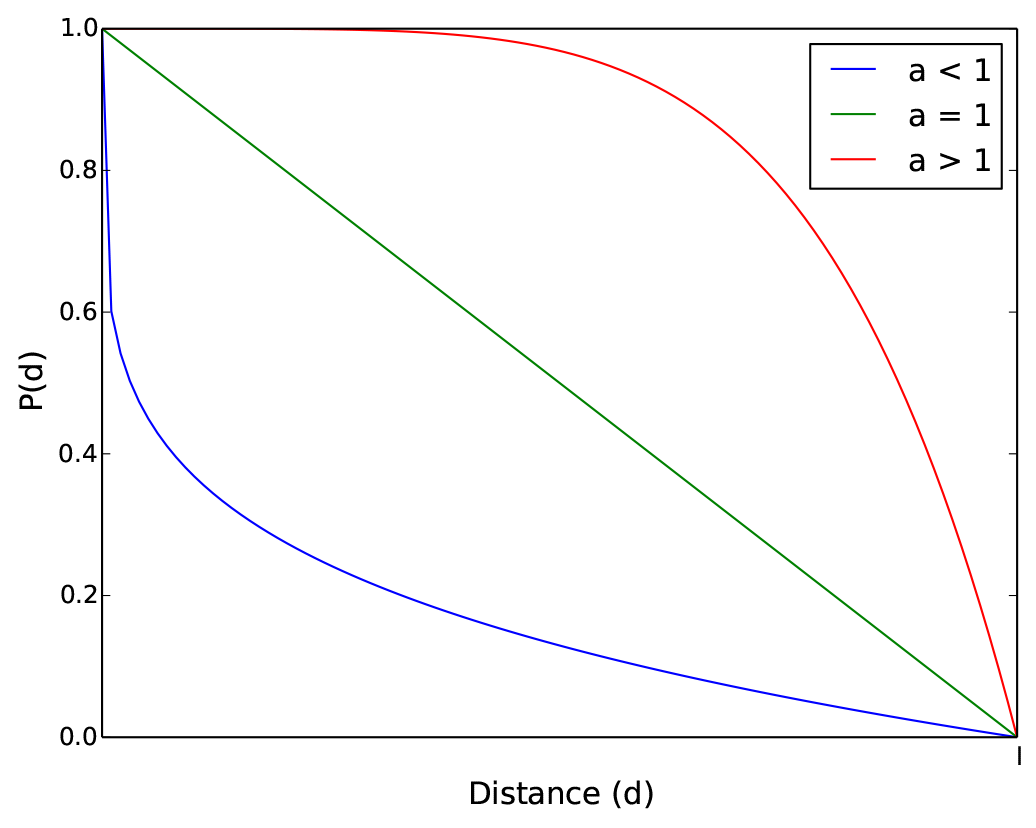
The probability function *P* (*d*) (cf. Eq. 3) is convex for 0 < *a* < 1 (blue), linear for *a* = 1 (green) and concave for *a* > 1 (red).

### Parameterizing the probability function

Feasible combinations of parameters were selected using a brute-force method. Novkovic et al. reported a representative FRC network to contain 176 nodes and 685 edges [9]. These measurements were used as the desired size of our simulated networks. 176 nodes were placed randomly in the volume and connected through the probability function with a range of values for the two parameters *a* and *l* in the function. For each combination of parameters the expected number of edges in the generated network was calculated:

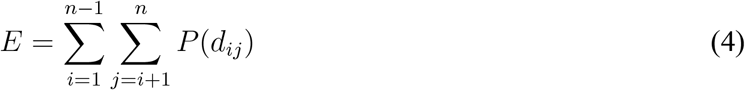

where n is the number of nodes, *d_ij_* is the distance between nodes *i* and *j* and *P* (*d_ij_*) is the probability function. The equation gives the expected number of total edges in the graph by summing the probability for each possible connection between any of the nodes in the graph. Whenever the expected number of edges was within 10 percent of the desired number of edges, e = 685 ± 68, the values of the parameters of *P* (*d*) were saved, and the corresponding values for *σ* and *ω* were calculated.

## Results

### Small-world properties emerge in a wide range of networks

The probability function *P* (*d*) (Eq. 3) defines the likelihood that two nodes at distance *d* are connected by an edge of the network. A range of values for maximum connection length *l* and shape parameter *a* were tested (0 < *a* < 5, 0.1 < *l* < 0.5), i.e. 50 random values were chosen uniformly within these ranges for both parameters, and for every possible combination of those values the expected amount of edges was calculated using Eq. 4. For each of the resulting networks with the “correct” number of edges, 685 ±86, small-world metrics were calculated (Fig. 2). Combinations of parameters that resulted in the desired amount of expected edges existed for *a* < 1, *a* = 1 and *a* > 1 (Fig. 2A), corresponding to convex, linear and concave functions, respectively (Fig. 1). Selecting networks with the correct number of edges, 685 ± 68, values within the range 0.1 < *l* < 0.5 result in *σ* < 1 and −0.6 < *ω* < 0.6 (Fig. 2B).

**Figure 2:**
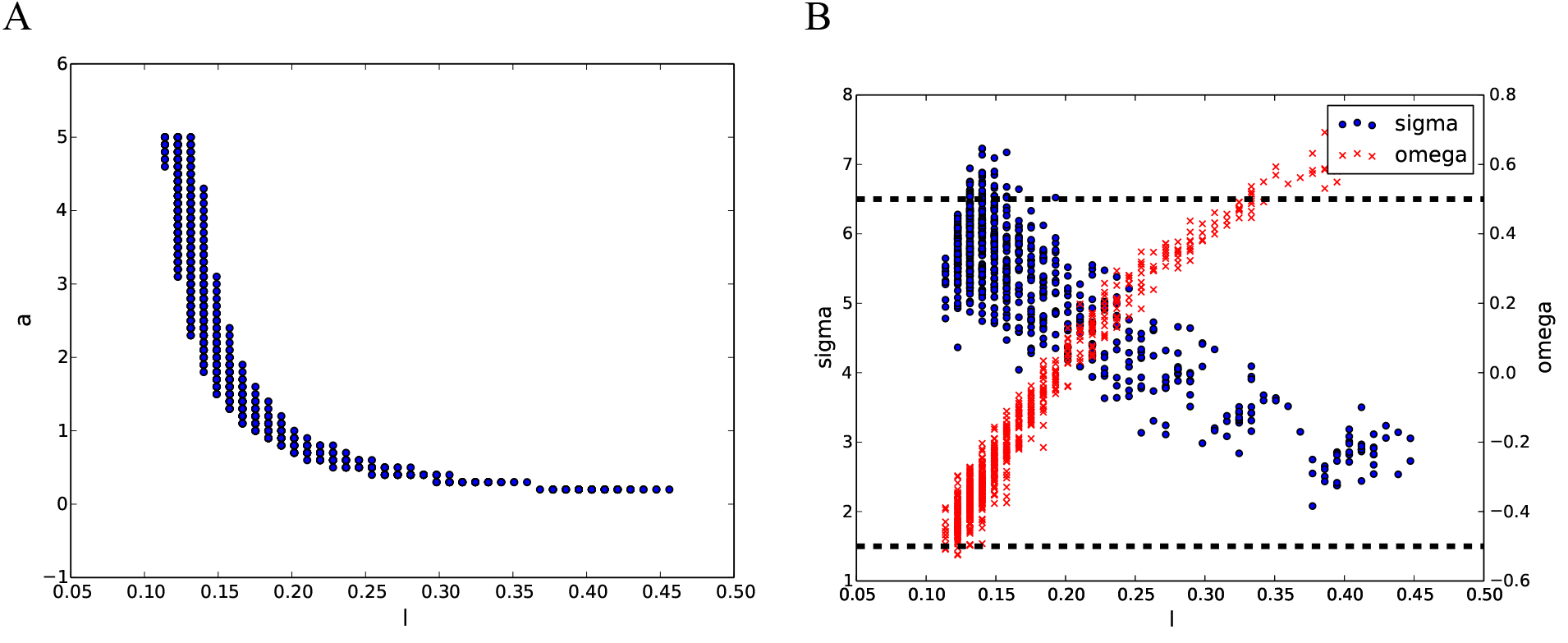
Parameter ranges of the probability function P(d) with the desired amount of edges and corresponding small-world metrics. Panel (A): values for the shape parameter, a, and the maximal distance, l, resulting in 685 ± 68 expected edges for n = 176. Panel (B): values for *σ* and *ω* obtained with each combination of a and l from Panel (A). Results are based on 10 networks for each combination of a and l.

This indicates that small-world properties are to be expected for networks within this range, except for the extremes of the range where *ω* < − 0.5 or *ω* > 0.5 respectively. Notably, *ω* is closest to 0, and thus small-world effects are strongest, around l = 0.2 and a = 1, corresponding to a simple linear probability function. Lattice-like properties (indicated by *ω* < 0) are observed for low values of *l* and when *a* > 1, corresponding to a concave function.

As *a* → ∞, the probability function approaches a step function where the probability of connecting is 1 if the distance *d* is smaller than the threshold *l*, and *P* (*d*) = 0 otherwise. This will generate a structure that is very similar to a regular 3D lattice. For the step function, the desired amount of edges was obtained for *l* = 0.1, resulting in networks with *σ* = 6.0 and ω = −0.5, which is still within the range to be classified as small-world, albeit on the edge. These results indicate that small-world properties naturally emerge in a variety of networks generated through a wide variety of geographical preferential attachment rules, without the requirement for extremely specific parameters, or additional alterations.

### Simulating the observed FRC network

The representative FRC network presented by Novkovic et al. contains 176 nodes and 685 edges, and had *σ* = 6.7 and *ω* = −0.27, indicating small-worldness with lattice-like properties [9]. The maximal distance between intersections of FRC strands is about 37 μm (26), suggesting a maximal connection length of 75 μm (which is scaled in the model to *l* = 0.25). Setting *l* = 0.25 and *a* = 1 in Eq. 3 generates networks with *σ* ≈ 5 and *ω* ≈ 0.25, which is small-world, but not lattice-like. In order to approximate the observed FRC network more accurately, we opted for a slightly more complicated distance function, i.e. a Hill function with a threshold for the maximal connection length:

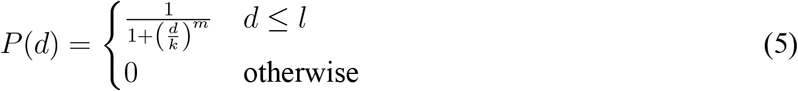

where *k* is the distance for which *P* (*d*) is half maximal, *m* determines the steepness of the function and *l* is the threshold for the maximal length. This Hill function results in sigmoid decay of the probability to form a connection as distance increases, resulting in a high ratio of short to long connections. For *l* = 0.25, *k* = 0.1 and *m* = 7, the generated networks have *σ* ≈ 6 and *ω* ≈ − 0.3 (Fig. 3B), which closely approaches the values found in the FRC network by Novkovic et al [9] (Fig. 3A). Equally, both the degree distribution (Fig. 3C) and the edge length distribution (Fig. 3D) of the generated networks are in accordance with the observed properties of the FRC network [9]. In order to test the robustness of the model, simulations were also run with *l* = 0.17 and *l* = 0.34, corresponding to a length threshold of 50 μm and 100 μm respectively. Similar values of *σ* and *ω* for all three values of *l* for somewhat different combinations of *m* and *k* (see Fig. S1). This indicates that results do not depend on the maximal connection length. These results show that geographical preferential attachment is sufficient to generate a topologically realistic FRC network.

**Figure 3:**
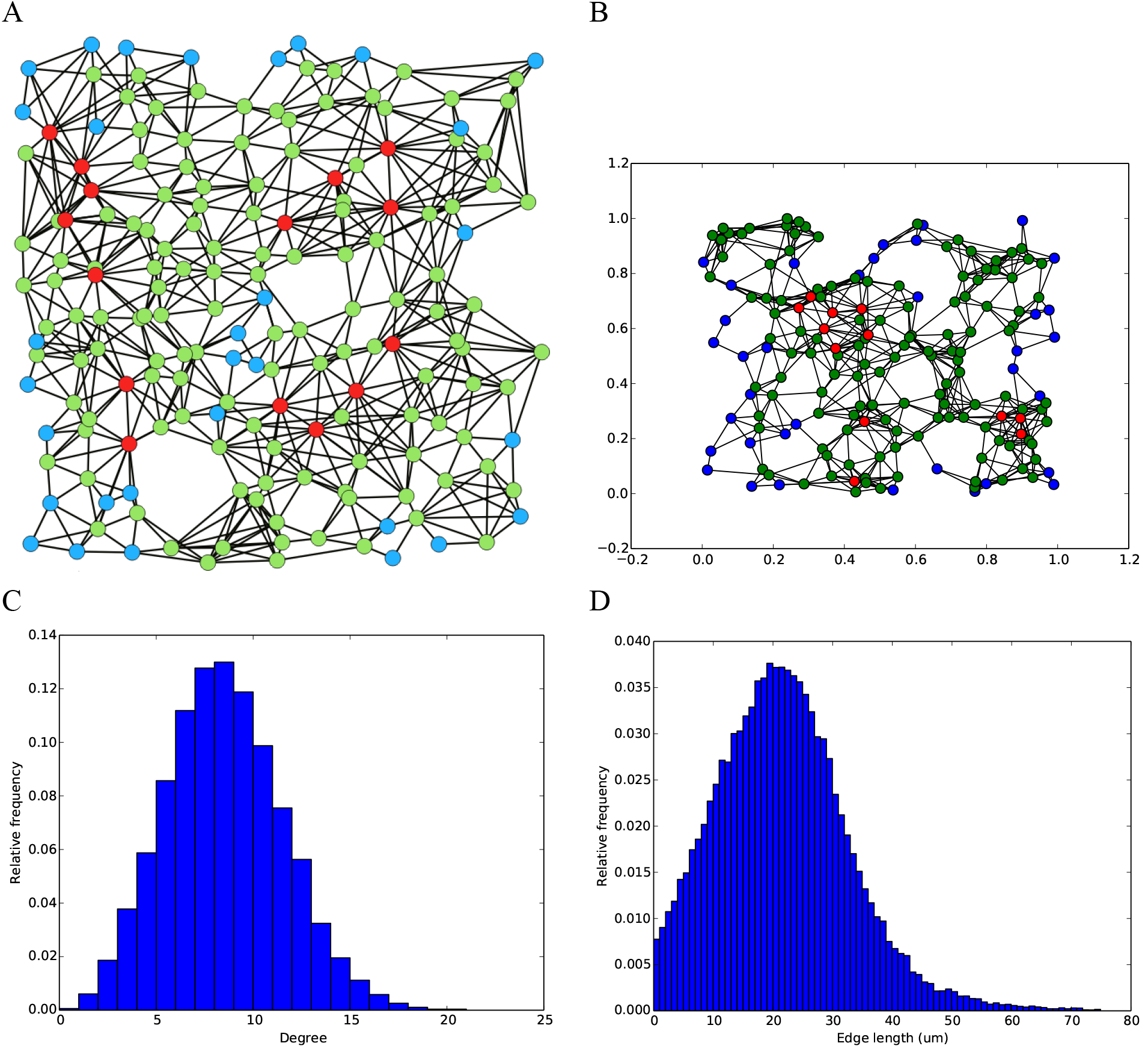
Comparison of an observed and a simulated FRC network. Green nodes have 5 or fewer edges, blue nodes have 6 to 11 edges and red nodes have 12 or more edges. (A): The FRC network depicted in Figure 1d of Novkovic et al. (8), with n = 176, e = 685, *σ* = 6.70 and *ω* = −.27. (B): a representative simulated network with n = 176, e = 677, *σ* = 6.07 and *ω* = −.28. (C): degree distribution of the generated networks. (D): edge length distribution the generated networks. Distributions based on 100 generated networks.

### Influence of network size

Current confocal microscopy techniques are incapable of generating high-resolution 3D images of the FRC network, limiting observations to thin cross-sections [9]. Using the model developed above, it is possible to simulate a realistic 3D FRC network. In order to investigate the influence of the thickness of the cross-section on network properties, the size of the grid was gradually expanded from 1x1x0.1 to 1x1x1 (corresponding to a network of 300x300x300 μm), while keeping the node density constant.

Increasing the thickness of the slice was found to affect the average shortest path length and average degree (Fig. 4A), but not the small-worldness (Fig. 4B). *σ* increases almost fourfold as the network is expanded. This is in agreement with the observation that *σ* scales with the network size [24]. *ω* approaches 0 but remains negative, indicating that both the small-worldness and lattice-like properties of the network are preserved. The average degree of the network increased from 4 to 6 as thickness increased from 0.1 to 0.2 and continued to increase when thicker 3D networks were constructed, which is a natural result because the impact of missed connections at the boundaries decreases when one considers thicker slices. As a result of these additional connections, the average shortest path length initially decreases as the network size is increased. As network size increases further this effect saturates and the average shortest path length starts increasing again.

**Figure 4:**
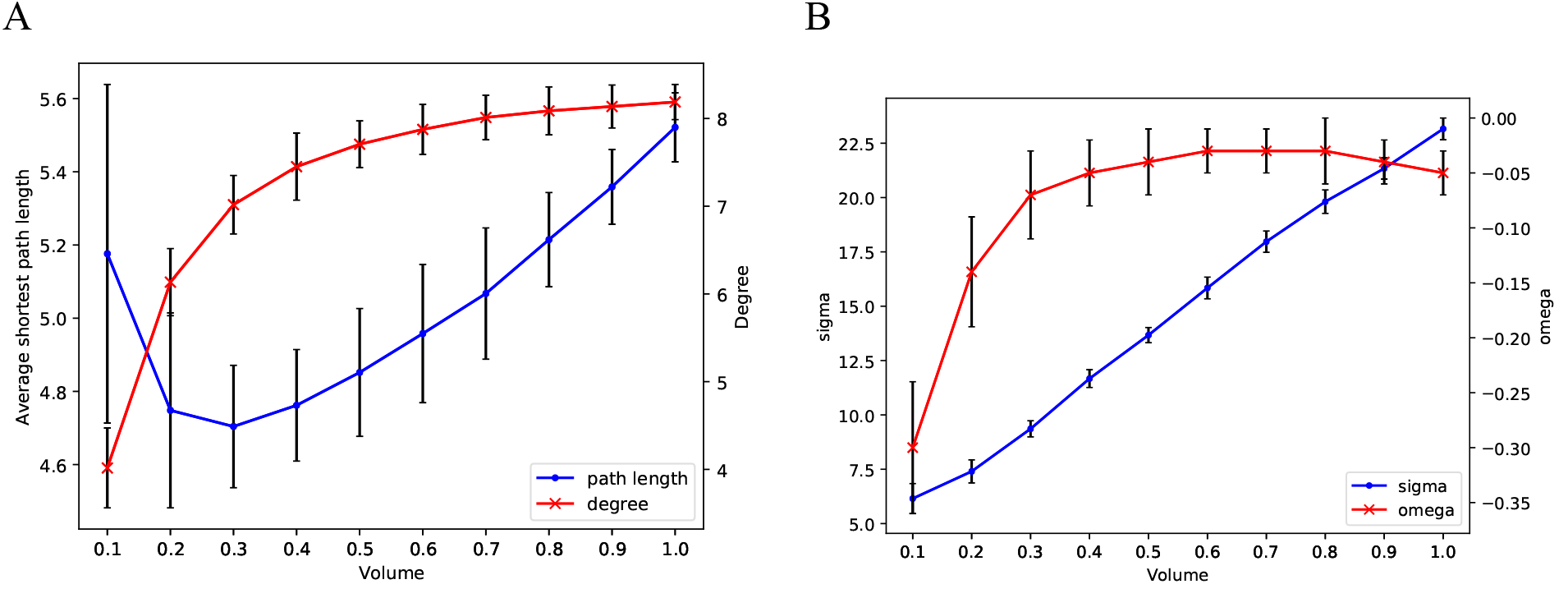
Increasing the thickness of the slice from 30 μm (volume 0.1×1×1) to 300 μm (volume 1×1×1). (A): Average shortest path length and average degree. (B): small-world metrics *σ* and *ω*. Bars indicate full range of values found for 50 independent simulations per volume size.

Subsequently, the effect of the size of the network on small-world properties was examined by increasing the volume of the cube in which the networks are generated. Network size was increased tenfold from 2.7×107 μm^3^ to 2.7×108 μm^3^, while node density and parameters of the probability function were kept equal (Fig. 5). Similar to the effect of increasing thickness, *σ* was found to increase further as network size increased. *ω* decreased as network size increased, indicating that the network becomes increasingly lattice-like. In Table S1 we provide all parameters of the generated networks.

**Figure 5:**
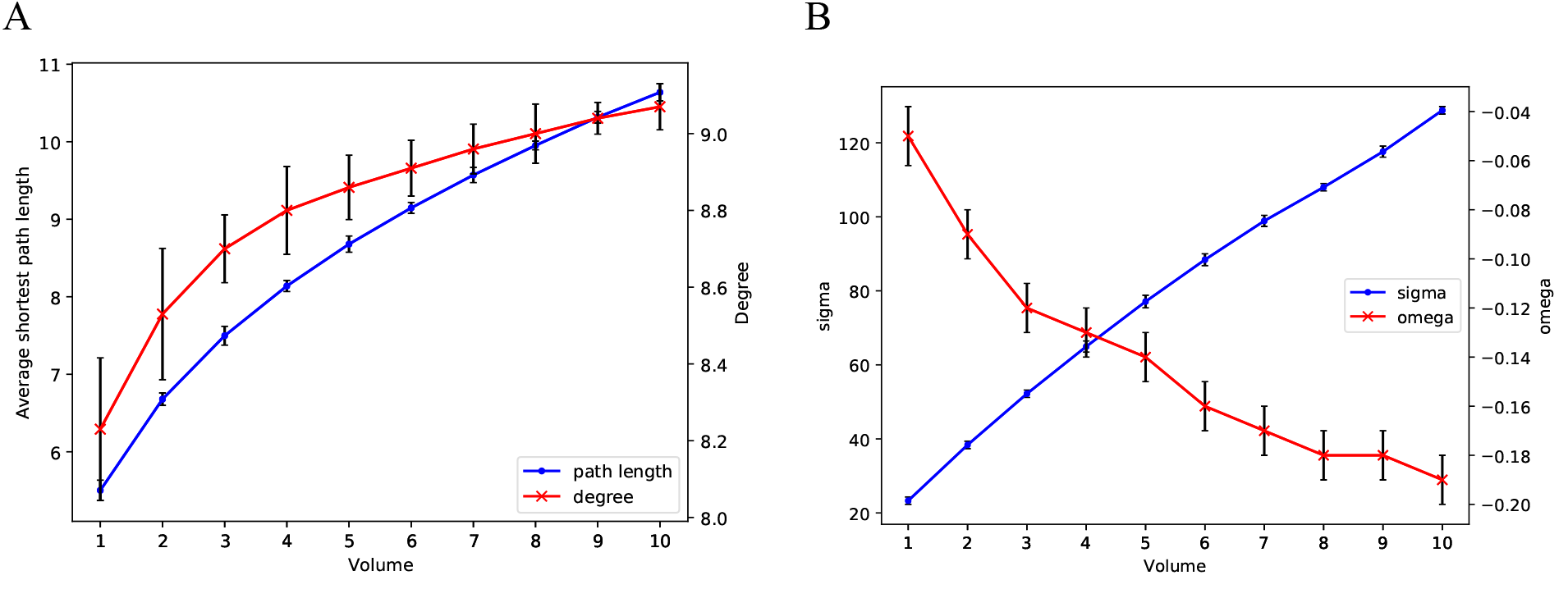
Increasing volume from 1 (2.7*107 μm^3^) to 10 (2.7*108 μm^3^). (A): Average shortest path length and average degree. (B): small-world metrics *σ* and *ω*. Error bars indicate range of values found for 50 independent simulations per volume size.

The average shortest path length of the observed networks increases much faster than the shortest path length of an equivalent random network (LR), whereas the clustering coefficients of the networks converge to the values = 0.38, CL = 0.71 and CR = 0.0001 (Table S1B). Thus, when the network becomes infinitely large, Eq. 2 for *ω* reveals that 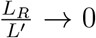, and hence that 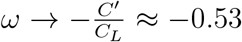, remaining on the edge of the range in which networks are classified as small-world. It should be noted that even though these networks are considered small-world according to *σ* and *ω*, they do not adhere to the logarithmic scaling between the number of nodes in the network and average shortest path length (which is impossible due to the presence of a maximal connection length threshold forbidding long connections).

### Shortest path length comparison

Next, we examined the average shortest path length of the generated networks. As lymphocytes follow the edges of the FRC network to migrate through the lymph node (26), the average shortest path length gives insight into the distance lymphocytes need to travel in order to traverse the network. Average shortest paths were measured in generated FRC networks for various network sizes and compared to random networks with an equal number of nodes and edges, and to complete networks with an equal amount of nodes that were fully interconnected. Both topological and physical average shortest path lengths were studied by measuring geodesic distance (Fig. 6A) and Euclidean distance (Fig. 6B), respectively.

**Figure 6:**
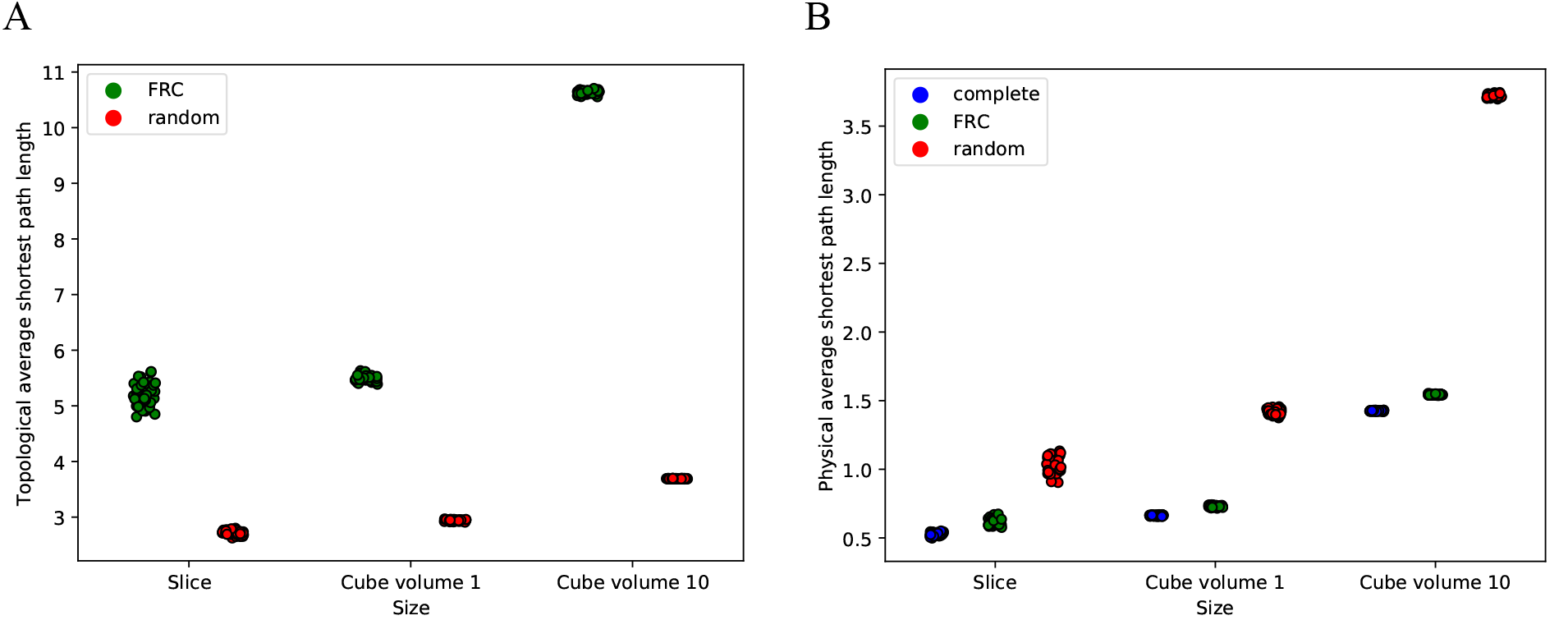
Physical and topological shortest path length comparison. (A): Topological average shortest path length (geodesic distance) of the generated FRC networks and equivalent random networks. (B): Physical average shortest path length (Euclidean distance) of the generated FRC networks, equivalent random networks and equivalent complete networks. Network sizes are 1x1x0.1, 1x1x1 and 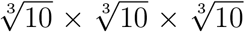 for slice, cube volume 1 and cube volume 10 respectively. Results are based on 50 independent simulations for each size and network type.

FRC networks were found to have a higher topological average shortest path length than equivalent random networks, indicating that on average more nodes need to be traversed in the FRC network in order to reach another node. This is caused by the fact it is more difficult (or even impossible) to form long connections in the simulated FRC networks, requiring more connections to connect nodes that are physically far apart. The topological average shortest path length of the FRC networks increases strongly as network size increases.

However, the physical average shortest path length is found to be much shorter in the FRC networks than random networks, indicating that the path taken in the FRC network is much closer to a straight line. In fact, the physical average shortest path length of the FRC network only slightly exceeds the path length in an equivalent complete network for all network sizes (Fig. 6B). This shows that the structure of the FRC network allows for paths between nodes that are nearly as short as possible, while requiring only a fraction of the connections. These relatively straight paths are likely the result of the lattice-like properties of the FRC networks.

### The FRC network closely resembles a regular lattice network

As the size of the FRC network is increased, its properties become increasingly lattice-like. The average shortest path length increases quickly due to the maximum length threshold, while the high clustering coefficient is retained. For an infinite network size, *ω* converges to −.53, which is on the border of being small-world. These observations raise the question whether the used small-world metrics adequately describe the small-worldness of the network. We therefore compared the clustering coefficients and average shortest path lengths of the FRC networks to those of regular lattice networks of the same dimension and size (Fig 7A).

**Figure 7:**
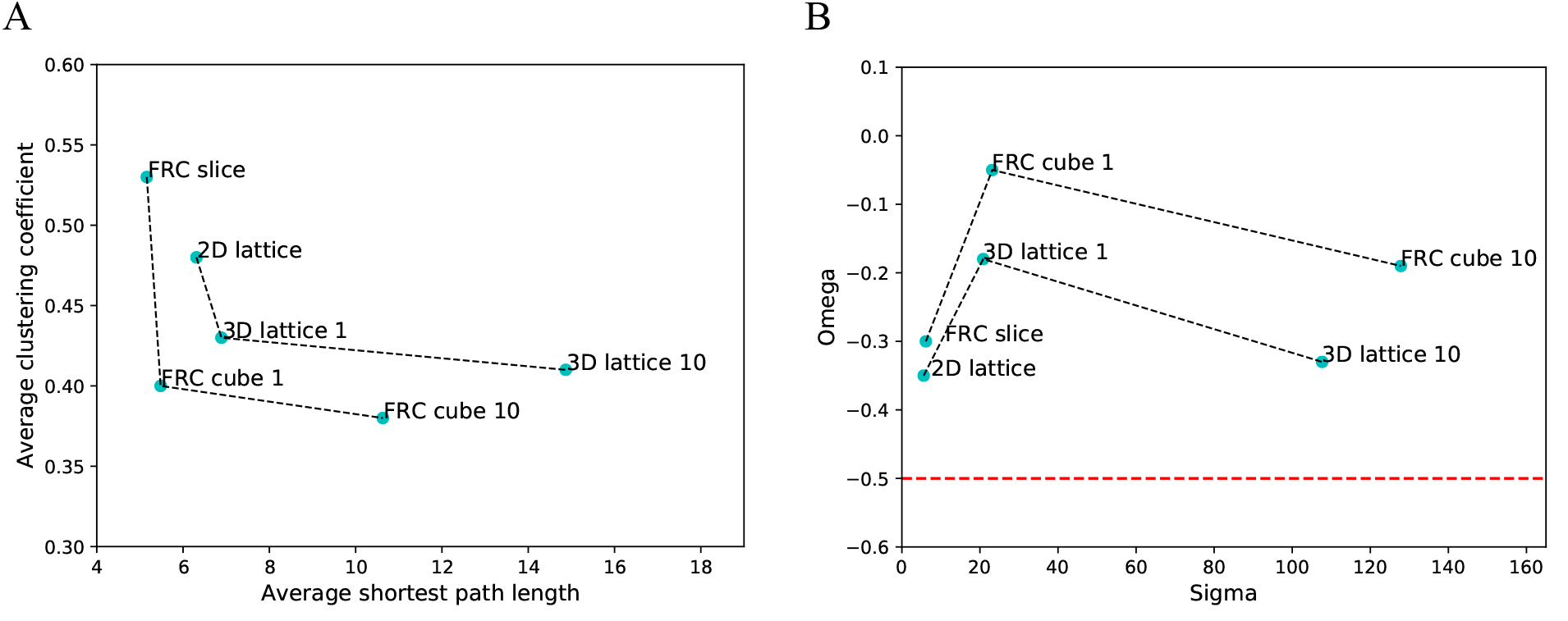
Comparison of FRC networks to corresponding lattices. (A): Average clustering coefficient and average shortest path lengths. (B): *σ* and *ω* values, with the red line indicating the small-world threshold. The lattice networks that we use are 8-regular 13x14 (n = 182, e = 649), 18-regular 12x12x12 (n = 1728, e = 13464) and 18-regular 26x26x26 (n = 17576, e = 210700), which corresponds to the FRC slice (n = 176, e = 182), cube volume 1 (n = 1760, e = 14450) and cube volume 10 (n = 17600, e = 159632), respectively. The lattice networks do not have wrapped boundary conditions. Dashed lines connect the different sizes of each type of network.

The properties of the FRC networks resemble those of “equivalent” lattice networks for all sizes. This indicates that the FRC network can approximately be described as a 3D lattice, rather than having a specific small-world structure. Indeed, the small-world metrics for these lattice networks fall within the range that indicates small-worldness (*σ* > 1, –0.5 < *ω* < 0.5), showing that a lattice-like structure is sufficient to be classified as small-world (Fig 7B).

## Discussion

We studied topological properties of general cellular networks and compared them to the FRC network, both in thin slices and networks expanded to symmetric 3D cubes. This has provided two new insights in the formation and topological properties of the FRC network.

First, results of the network simulations show that small-world properties are likely to emerge naturally in cellular networks with physical connections. Various different probability functions resulted in small-world networks for a range of parameter values. This indicates that small-worldness as measured by the *σ* and *ω* parameters is a robust and naturally occurring property of these networks, and need not require evolutionary pressure [9].

Second, simulation of the FRC network with accurate topological properties and its subsequent expansion to 3D networks showed a significant increase in small-worldness and a doubling of the average degree in the network. Increasing the network size further resulted in higher lattice-likeness of the network and a slight increase in average degree. This indicates that not all network properties are correctly inferred from thin slices. The FRC network is a 3D structure in the lymph node, reaching up to several millimeters in diameter [3]. Calculating network properties from a thin slice of the network is expected to underestimate the average degree in the network, and to provide inaccurate topological properties.

The observation that small-world properties emerge naturally and persist even for larger network sizes raises questions on the accuracy of the *σ* and *ω* metrics. Expansion of the network size showed that small-worldness was conserved in larger networks, and values for *σ* and *ω* were found to remain in the small-world range even when the network becomes very large. However, the average path length of the simulated FRC networks grows much faster than that of equivalent random networks. Due to the presence of a maximal connection length, the average path length of the FRC network is proportional to the network size, rather than having the logarithmic proportionality that would be expected in a “true” small-world network. As the network size increases and the average shortest path length becomes much larger than that of a random network, *ω* is reduced to a ratio of the clustering coefficients of the FRC network and its equivalent lattice network. The average clustering coefficient of a network is a local property of the nodes in the network and converges as the network size is increased. The FRC network is sufficiently clustered that the ratio of the clustering coefficients remains on the verge of being classified as a small-world network. In the original definition by Watts and Strogatz, small-worldness requires the path length to be similar to that of an equivalent random network [20]. This is clearly not the case for the large FRC network, and illustrates that the small-world metrics *σ* and *ω* are not always accurately assessing small-worldness.

Comparing simulated FRC networks with lattice networks of equivalent size and dimension shows a close resemblance between the types of networks, both in network measures as well as small-world metrics. Therefore, it seems likely that the observed small-world metrics are not the result of a specific small-world structure present in the FRC network, but are rather due to its lattice-like structure which happens to fall within the small-world range of the *σ* and *ω* metrics. This could indicate that the observed small-world topology does not play an important role in the functioning of the FRC network. Lymphocytes are known to follow the FRC network during migration [27, 28], but this does not require a small-world structure to be present. Indeed, the relatively small physical shortest path length observed in the network is caused by the lattice-like properties. Thus, FRC networks may have small-world properties just because they are formed by preferential attachment to nearby neighbors.

## Acknowledgement

JT was supported by the Dutch Cancer Society – Alpe d’HuZes foundation (project 10620).

